# The design and implementation of restraint devices for the injection of pathogenic microorganisms into *Galleria mellonella*

**DOI:** 10.1101/2020.03.10.985481

**Authors:** Lance R. Fredericks, Cooper R. Roslund, Mark D. Lee, Angela M. Crabtree, Peter B. Allen, Paul A. Rowley

**Affiliations:** Department of Biological Sciences, University of Idaho, Moscow, ID 83844, USA; Department of Chemistry, University of Idaho, Moscow, ID 83844, USA

## Abstract

The injection of laboratory animals with pathogenic microorganisms poses a significant safety risk because of the potential for injury by accidental needlestick. This is especially true for researchers using invertebrate models of disease due to the small size of the animals and the required precision and accuracy of the injection. The immobilization of the greater wax moth larvae (*Galleria mellonella*) is often achieved by grasping a larva firmly between finger and thumb. Needle resistant gloves or forceps can be used to reduce the risk of a needlestick but can result in animal injury, a loss of throughput, and inconsistencies in experimental data. Immobilization devices are commonly used for the manipulation of small mammals, and in this manuscript, we describe the construction of injection chambers that can be used to entrap and restrain *G. mellonella* larvae prior to injection with pathogenic microbes. These devices significantly reduce the manual handling of larvae and provide an engineering control to protection against accidental needlestick injury, while maintaining a high rate of injection.

## Introduction

The larvae of the greater wax moth *Galleria mellonella* is an important animal model for studying host-pathogen interactions and for the discovery of novel antimicrobial therapeutics. The popularity of this model organism is driven by the low cost of purchase and the reduced ethical concerns for the experimental manipulation of insects. This allows the challenge of a large number of larvae in a single experiment, which can improve the statistical power of an assay. Starting in the 1940s, a diversity of viral, bacterial, fungal, and nematode pathogens, have been studied for their ability to cause disease in *G. mellonella* larvae [1–15]. Importantly, *G. mellonella* can be maintained at mammalian body temperature and the outcomes of infection can reproduce that of mammalian animal models [16–18]. This is likely due to similarities in the innate immune response to pathogens mediated by elements of cellular and humoral immunity between insects and mammals [19–21]. There are several methods to initiate an infection of a larva with a pathogen, including topical application, feeding, oral gavage, and direct injection into the hemocoel. The latter method is favored because of the ability to control the timing of infection and the dosage of pathogen.

Despite the benefits of *G. mellonella* larvae, there are challenges with using this animal model, most notably standardizing the health and developmental stage of the larvae. This is especially challenging when they are purchased from commercial sources that are primarily focused on providing feed for the pet and angling communities [22,23], although there are efforts to develop commercial pipelines for scientific-grade larvae [24]. Another difficulty is the manipulation and restraint of small larvae during experimental injection of pathogenic microbes. These experiments require both biological containment and adequate protection of personnel from needlestick injuries and accidental infection with a pathogen. The most basic technique of larval injection calls for the immobilization of a larva between finger and thumb during the injection process [9,11,14,25]. With the operator’s hands protected by latex gloves, this method offers the maximum dexterity to enable the immobilization of a larva prior to injection. However, the proximity of a pathogen-loaded needle to inadequately protected fingers presents a significant biological safety hazard that exposes personnel to a high risk of accidental needlestick injury and infection by pathogenic microbes. This immobilization procedure is in conflict with biosafety guidelines for the implementation of policies for improved work practices that minimize needlestick injuries whenever possible [26]. The safe handling of *G. mellonella* larvae can be achieved with the use of needle resistant gloves or forceps, but with a loss of manual dexterity and the potential to cause the animal injury or stress that can alter their susceptibility to a pathogen [27]. Furthermore, needle-resistant gloves are generally made of porous material that requires covering with a pair of disposable gloves to prevent biological contamination, which further limits manual dexterity. Humane and safe physical restraint devices have been used routinely for the safe handling and manipulation of many different types of laboratory animals [28], and an immobilization device has been previously developed for *G. mellonella* larvae. The “Galleria grabber” entraps larvae between layers of a sponge prior to injection [29]. However, the reuse of the sponge for multiple injections increases the chance of contamination by pathogenic microbes or infected hemolymph and requires frequent decontamination to minimize cross contamination and maximize biological containment. In this study we present two simple restraint devices that can be made from a variety of laboratory consumables or fabricated from acrylic glass (also known as poly(methyl methacrylate) or Plexiglass). These devices can be used to restrain *G. mellonella* larvae in preparation for injection. The protocol is easy to perform, significantly reduces the manual handling of larvae, maintains injection speed, and provides increased protection of the operator from accidental needlestick injury and laboratory-acquired infection.

## Methods

### Strains, species and culturing of yeasts

*C. glabrata* ATCC2001 was maintained using yeast extract peptone dextrose growth media (YPD). Prior to injection, yeasts were grown overnight at room temperature to stationary phase in a 2 mL culture of liquid YPD medium. Stationary phase cultures were diluted 1/20 into a 125 mL flask and grown at room temperature until an OD_600_ of 1.5 was reached. Hemocytometer counts of these cultures were used to determine the number of yeasts used for each injection. Prior to injection, yeast cells were harvested by centrifugation and suspended in PBS.

### *G. mellonella* larvae handling, care, and disposal

Larvae were ordered from www.premiumcrickets.com using the “weather protect” service to maintain the temperature of the larvae during overnight shipping. Upon arrival the larvae were stored in the dark in wood shavings at 17°C and allowed to acclimatize for at least 2 days to control for the adverse physiological consequences of shipping [27]. *G. mellonella* larvae were used within 1 week due to the known physiological consequences of long-term storage of larvae [30]. Healthy *G. mellonella* larvae were selected by weight (175-225 mg), uniformity in color (little to no melanization), and responsiveness to touch. Prior to injection, larvae were incubated at 37°C for 16 hours to allow for acclimatization to the assay temperature. Dead or unhealthy larva that are observed after the pre-incubation period are removed from the study prior to injection. To safely dispose of larvae exposed to pathogenic microorganisms, larvae were placed in secondary containment and incubated at −20°C for 16 hours before sterilization by autoclaving and disposal.

### Fabrication of restraint devices for *G. mellonella*

The primary consumable used for the creation of the restrain device was the 250 μL VistaLab micropipette tip (catalog number: 4058-2000). Other brands of pipette tips have been tested for their compatibility with this method (S1 Table). Micropipette tips were cut with a sterilized razorblade at predefined points to enable assembly (Fig 1). To construct the restraint device fabricated from acrylic glass, transparent clear acrylic Plexiglass was purchased from AliExpress (1 mm × 100 mm × 100 mm) and was cut to the desired shape with a CO_2_ laser cutter (BOSS Laser LS-1416) using software provided by the manufacturer (available online in SVG format at thingiverse.com design ID 4170068, “LarvaWormCorralV4”).

**Fig 1.**
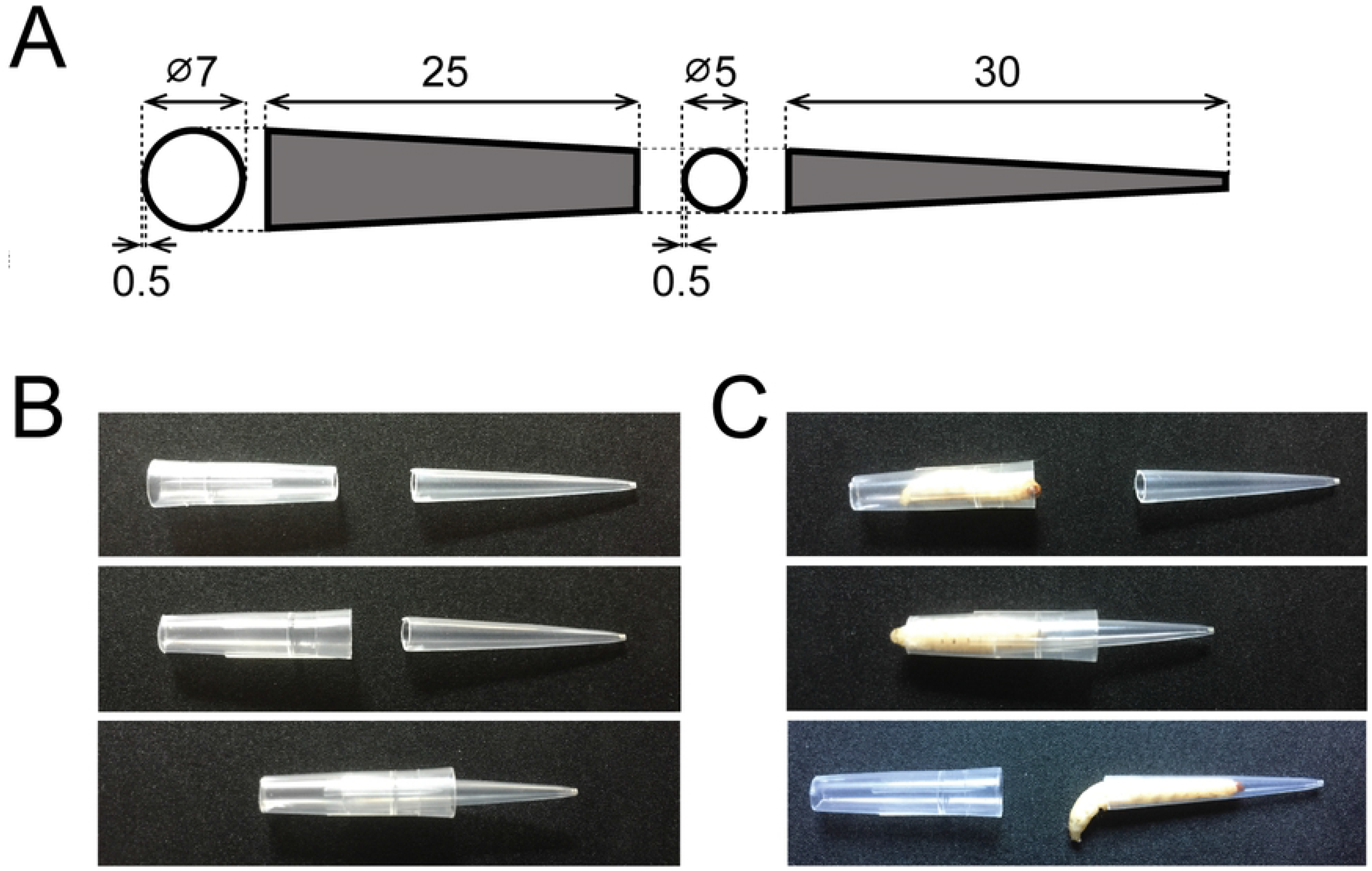
Entrapment of *G. mellonella* in an injection chamber constructed from a disposable micropipette tip. (A) Dimensions of a 250 μL VistaLab pipette tip indicating the cut site to create two 5 mm openings. (B) Assembly of the injection chamber without larvae (C) Top: Assembly of the injection chamber with a larva captured abdomen-first into the larger half of the chamber. Middle: A larva is entrapped by enclosing the chamber. Bottom: A larva is released by opening the chamber. Measurements are presented in millimeters.

### Injection of restrained *G. mellonella*

Single injections were performed with Hamilton 700 syringe, Model 701 N, Volume: 10 μL, Point Style: 2, Gauge: 26s. Multiple injections were done with a Hamilton 700 syringe, Model 1750 LTSN SYR, Volume: 500 μL, Point style: 4 Gauge: 26s. The repeating dispenser used was the Hamilton PB600-1 repeating dispenser. Ethanol diluted in water to a final concentration of 70% was used to decontaminate all syringes and needles prior to and during injection of *C. glabrata*. To decontaminate a syringe, each is suspended in a beaker containing 70% ethanol with 1.5 cm of the needle submerged in the ethanol bath and the syringe filled with ethanol for a total contact time of greater than 10 min. To increase throughput, either multiple Hamilton syringes or a repeat dispenser are used to ensure a continuous and rapid rate of injection. Prior to injection, ethanol is dispensed from the syringe and washed three times with sterile distilled water before loading with a suspension of yeast cells before injection. Entrapped larvae are presented for injection by holding the injection chamber between finger and thumb that are protected with a HexArmor PointGuard^®^ Ultra 4041 High Performance Needle-Resistant Search Gloves (Performance Fabrics Incorporated). A disposable laboratory glove is used to prevent biological contamination of the porous needle-resistant glove. Restraint devices can be decontaminated with 70% ethanol or 10% bleach after ever use to prevent contamination with pathogenic microbes. While it is possible to sterilize the micropipette devices using an autoclave, this method is not recommended for devices made of acrylic glass [31].

### Data analysis

The graphical representation of the average survival rates of *G. mellonella* larvae after injection and the Kaplan-Meier log-rank analysis were performed using R (version 1.1.419) with the packages “ggplot2”, “dplyr”, “survival”, and “survminer”. LT_50_ was calculated using the package “MASS”.

## Results and Discussion

### The restraint and injection of *Galleria mellonella* larvae using a reusable injection chamber

A central challenge to the manipulation and injection of *Galleria mellonella* larvae is the ability to restrain the animals without injury. Careless handling of a larva, often because of the use of needle-resistant gloves, can cause injury and result in premature death. To minimize these undesirable consequences, we have designed and tested two types of injection chamber for the restraint of individual *G. mellonella* larvae. These chambers can be constructed either from a disposable 250 μL or 1,000 μL micropipette tip of the required dimensions (Fig 1A and S1 Table) or assembled from laser cut acrylic sheet (Fig 2) and are designed to be fully reusable. To make the micropipette tip device, a tip is cut, and the two halves are used to entrap a larva with minimal handling (S1 Movie).

Specifically, a larva is captured within the wider end of the cut pipette tip and entrapped by inserting the second half of the pipette tip to seal the injection chamber at one end (Fig 1B). This procedure takes an average of 37 seconds (SD = 17 seconds). After entrapment, the escape time of larvae from the device was measured under ambient light conditions. After 30 and 60 minutes, 7% and 20% of larvae were observed to exit the device, respectively. This allows the loading of multiple larvae without the need for biological containment and allows a faster rate of injection due to the predictable presentation of the entrapped larva inside a chamber. Moreover, once entrapped, larvae generally wedge themselves into the chamber and remain motionless, even when turned to reveal their ventral side, which is likely due to their known aversion to light (Fig 1C and Fig 3) [32]. Larvae that are attempting to exit the device prior to injection can be persuaded to re-enter the chamber with a gentle touch to their abdomen. After injection, a larva can be released by gently pulling the two halves of the chamber apart to allow egress (Fig 1C). The design schematics for an acrylic glass injection chamber have been deposited in the Thingiverse public repository (https://www.thingiverse.com/), identifier number 4170068, LarvaWormCorralV4. An acrylic plastic sheet (1 mm thick) is cut to the designed dimensions with a CO_2_ laser cutter. Five layers are assembled as shown in Fig 2A. The three middle layers are cut with a triangular notch to hold the larva. To assemble, the layers of the device are stacked to construct a chamber, using a binder clip to secure them together (Fig 2B). Depending on the size of the larvae, the height of the chamber can be adjusted by removing or adding notched layers. Importantly, a chamber that is too large will enable the larva to turn in the device and not present its ventral side for injection. Larvae are coaxed into entering the chamber head-first by gently pushing their abdomen towards the chamber opening. Once loaded with a larva, the acrylic glass injection chamber has the same properties as the micropipette tip chamber with regards to the speed of injection.

**Fig 2.**
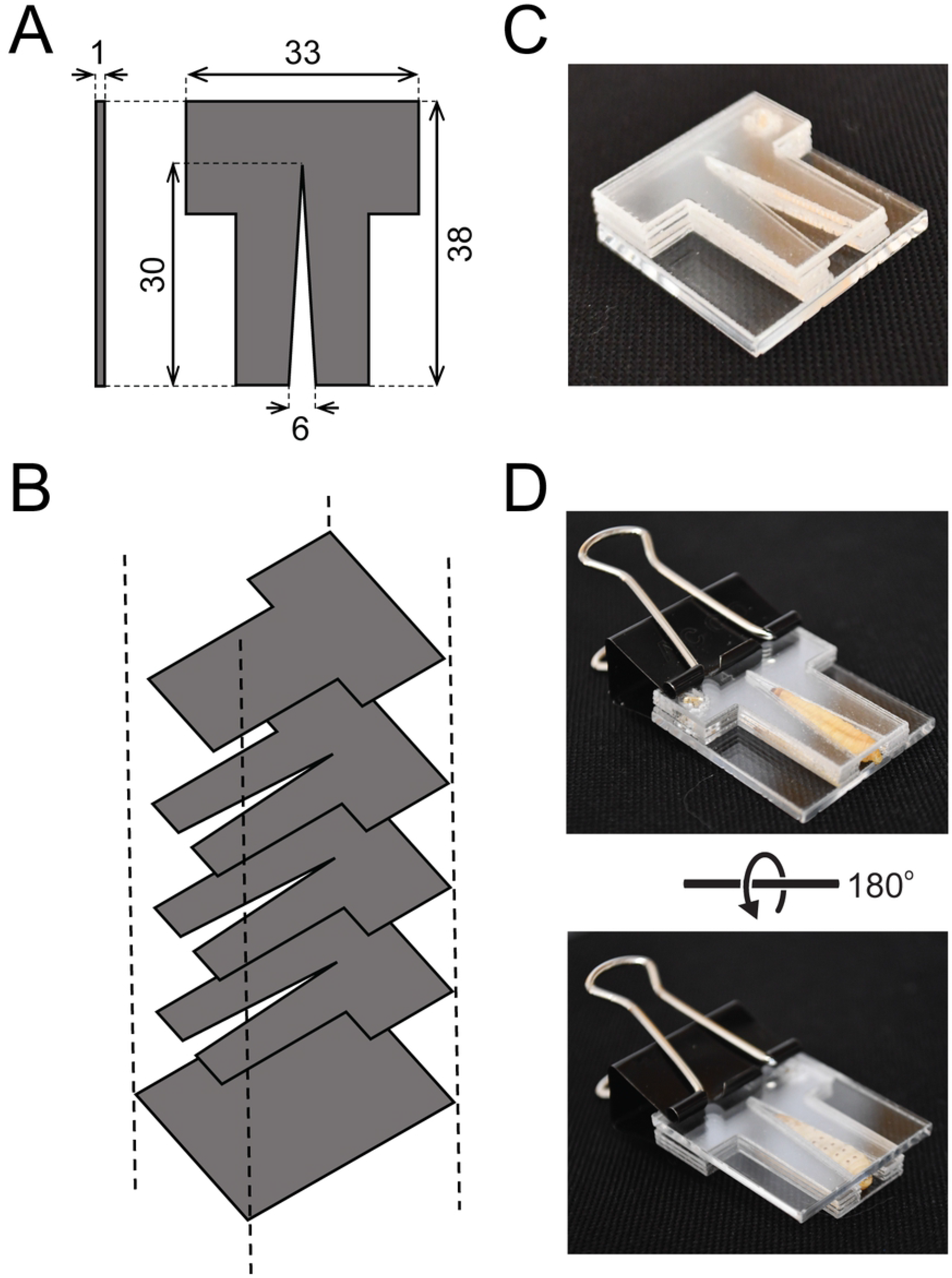
Entrapment of *G. mellonella* in an injection chamber constructed from laser cut acrylic sheet. (A) Dimensions of the injection chamber. (B) Schematic overview of the assembly of acrylic sheets without larvae. (C) Assembly of the injection chamber without a larva (D) Injection chamber with restrained larva captured head-first into the chamber. Ventral side is visible upon rotation of the chamber with the larvae unresponsive to inversion. Measurements are presented in millimeters.

To test the effectiveness of the micropipette tip injection chamber, 160 restrained larvae were injected with the pathogenic yeast *C. glabrata* or phosphate-buffered saline (PBS) (Fig 3). To inject a larva, a needle is inserted through the wide end of the injection chamber and used to pin down the larva at the last left proleg before puncturing the cuticle (S2 Movie). The needle is inserted along the long axis of the larva at a shallow angle of 10-20° beneath the cuticle to avoid puncturing the midgut. The needle penetrates to a depth of 5 mm, whereupon the plunger is depressed to eject the contents of the syringe into the hemocoel. The shallow angle and depth of injection can be verified as the needle is visible through the cuticle. In the rare case that an injected larva exudes excessive amounts of hemolymph or any white solids, then the individual is discarded because of the increased likelihood of severe injection-induced injury. After the withdrawal of the needle, the larva is placed in a 9 cm petri dish fitted with two 9 cm diameter disks of paper towel in the dark to promote the egress of the larva from the injection chamber. The injection time, using a repeat dispenser for a Hamilton syringe is 14 seconds per larva (SD +/− 2 seconds). The injection process with a single channel Hamilton syringe, which includes the filling of a single Hamilton syringe, injection, larval release, and decontamination of the needle and syringe, takes an average of 62 seconds per larva (SD +/− 12 seconds). Once injected, larvae were observed for 5 days to assess the physiological consequences of the injected microorganisms or PBS (Fig 4). Of the 63 larvae injected with PBS in three independent experiments, there was more than a 95% (60/63) survival after 5 days, with all larval death occurring within the first 24 hours after injection. Conversely, with the injection of different numbers of pathogenic *C. glabrata* (8 × 10^5^, 3 × 10^6^, and 5 × 10^6^), we observed dose dependent mortality and signs of infection, including melanization and reduction in larval movement before eventual death. Time to 50 % lethality (LT_50_) was calculated as 1.65 and 2.74 days upon injection of 3 × 10^6^, and 5 × 10^6^ *C. glabrata* cells, respectively. Lethality was greater than 80 % after 5 days when at least 3 × 10^6^ cells were injected. Conversely, 82 % of larvae survived the challenge with 8 × 10^5^ cells. Mortality caused by the injection of *C. glabrata* was significantly different than PBS upon the injection of 3 × 10^6^, and 5 × 10^6^ *C. glabrata* cells (p < 0.0001). These data are comparable to previous studies that used alternative injection methods to infect *G. mellonella* with *C. glabrata* ATCC 2001 [7].

**Fig 3.**
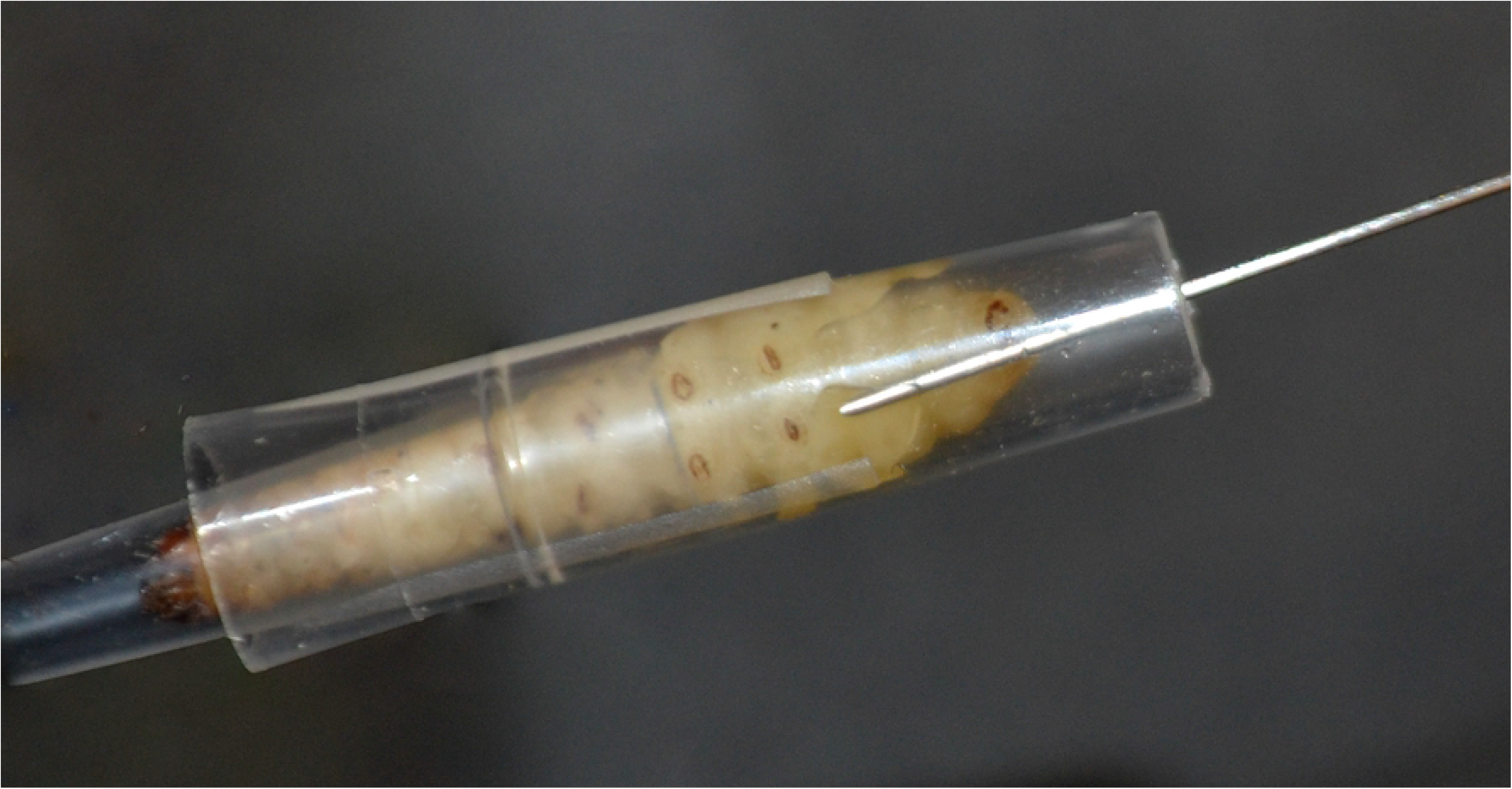
Ventral view of an entrapped *G. mellonella* during an injection of *C. glabrata* into the last proleg.

**Fig 4.**
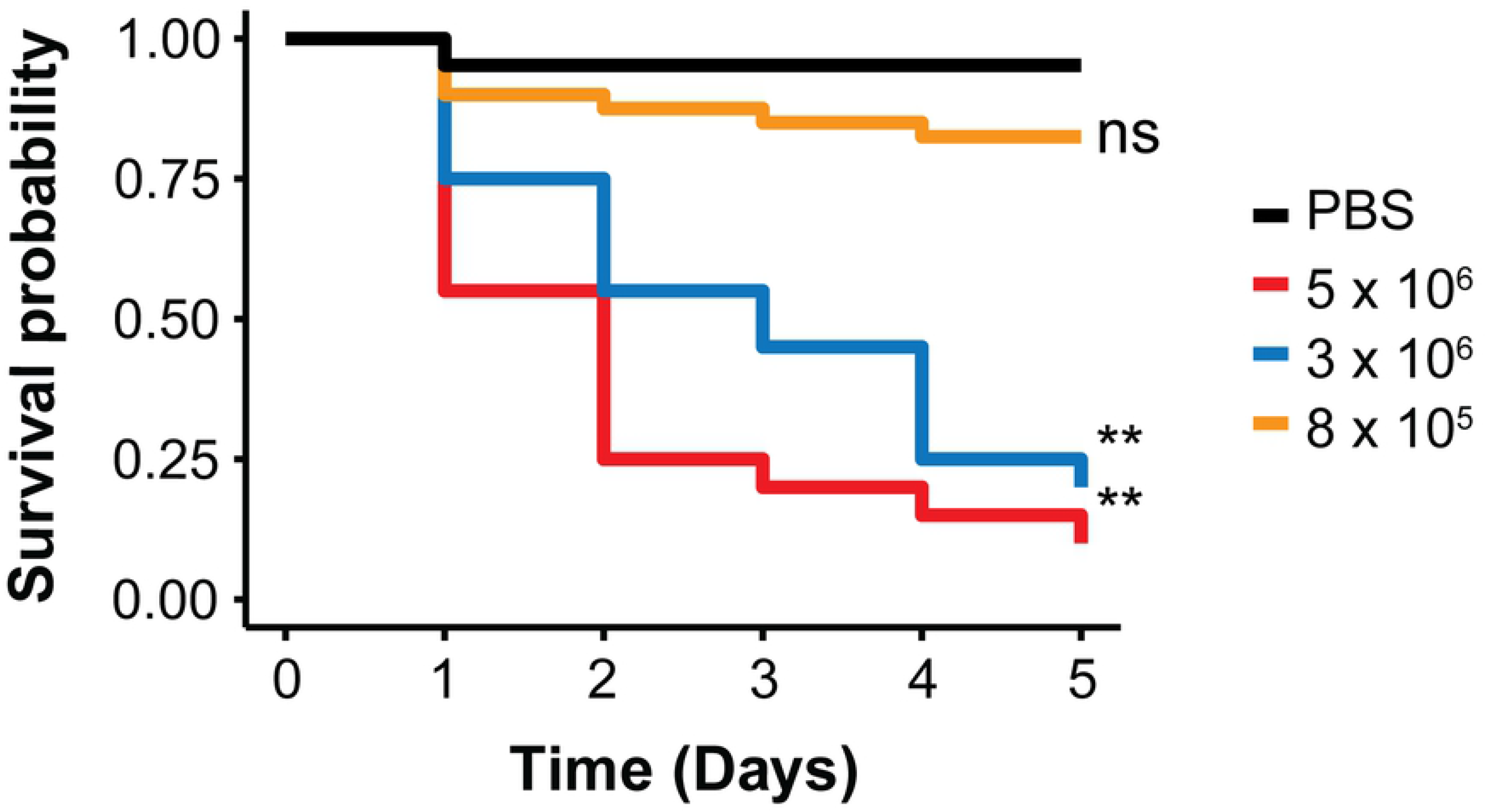
Survival rates of *G. mellonella* larvae over five days post-injection with three different inocula of *C. glabrata* cells or with PBS. Statistical significance was judged by the Kaplan-Meier log-rank analysis (** p < 0.0001, ns = not significant).

The method described in this report utilizing engineering controls, spatially separates the injecting needle and fingers, and when combined with a needle resistant glove, almost completely eliminates the risk of needle injury and the accidental infection of laboratory personnel with pathogenic microbes. The devices that we have described to restrain *G. mellonella* larvae are inexpensive, non-porous, and can be either discarded or reused after sterilization or decontamination. We have found that the ability to entrap multiple larvae before injection can enable a high rate of injection when using a repeat dispenser with a Hamilton syringe. Moreover, this protocol is most effective with two people working together, one loading larvae into injection chambers and the other performing the injections. The described method for the restraint of *G. mellonella* larvae offers a rapid and reproducible method for the injection of hazardous microorganisms.

## Funding

The research was supported by funds provided to PAR by the Institute for Modeling Collaboration and Innovation at the University of Idaho (NIH grant P20 GM104420), the Institutional Development Award (IDeA) from the National Institute of General Medical Sciences of the National Institutes of Health under Grant #P20GM103408, the National Science Foundation Cooperative Agreement No. 818368 and NSF Science and Technology Center on evolution in action, DBI-0939454. Funding was also provided by the Office of Undergraduate Research at the University of Idaho awarded to LRF and CRR. The funders had no role in study design, data collection and analysis, decision to publish, or preparation of the manuscript.

## Acknowledgements

We would like to thank the University of Idaho Biological Safety Officer Megan Grennille, and Nathan Taggart for critical reading of this manuscript. We would also like to acknowledge the contribution of Madison Chapman for technical assistance in the sorting and husbandry of *G. mellonella* larvae.

## Supporting information captions (if applicable)

**S1 Movie. Entrapment of a *G. mellonella* larva in an injection device constructed from a micropipette tip**.

**S2 Movie. Injection of a restrained *G. mellonella* larva in the last proleg with a Hamilton syringe**.

**S1 Table. Brand compatibility with injection chamber construction**

